# Improved estimation of phenotypic correlations using summary association statistics

**DOI:** 10.1101/2020.12.10.419325

**Authors:** Ting Li, Zheng Ning, Xia Shen

## Abstract

Estimating the phenotypic correlations between complex traits and diseases based on their genome-wide association summary statistics has been a useful technique in genetic epidemiology and statistical genetics inference. Two state-of-the-art strategies, Z-score correlation across null-effect SNPs and LD score regression intercept, were widely applied to estimate phenotypic correlations. Here, we propose an improved Z-score correlation strategy based on SNPs with low minor allele frequencies (MAFs), and show how this simple strategy can correct the bias generated by the current methods. Comparing to LDSC, the low-MAF estimator improves phenotypic correlation estimation thus is beneficial for methods and applications using phenotypic correlations inferred from summary association statistics.

## Introduction

Phenotypic correlation is an essential parameter that helps understand observational correlations between complex traits and the etiological perspectives underlying complex diseases. Conventionally, estimation of the phenotypic correlation between a pair of phenotypes, by definition, is straightforward in a sample where both phenotypes are measured. Depending on the distribution of each phenotype, the estimated phenotypic correlation serves as a sufficient statistic for many linear statistical models, such as ordinary linear and logistic regressions, allowing us to assess parameters such as odds ratios of risk factors on disease outcomes.

Since a large number of genome-wide association studies (GWAS) were conducted, many GWASed phenotypes had measurements in an overlapping set of individuals, where many were from more than one participating cohort in GWAS meta-analysis. In practice, inference of the phenotypic correlations across these phenotypes would be complicated if estimating using the conventional way, which requires individual-level phenotypic data and subsequent meta-analysis. Fortunately, the phenotypic correlations can be estimated based on established GWAS summary statistics, especially when the proportion of sample overlap between two GWASed phenotypes is large. Two state-of-the-art strategies were proposed:

1. *“Z-cut” estimator*. The phenotypic correlation can be estimated by the correlation between the two sets of GWAS estimated effects or Z-scores, assuming the genetic effect per SNP (single nucleotide polymorphism) is tiny or even null^1, 2, 3, 4^,
2. *LDSC intercept*. The phenotypic correlation can be estimated by the intercept of a bivariate linkage disequilib-rium score regression (LDSC)^5, 6, 7^.

Both estimators have reasonable performance in practice, however, bias exists for both strategies. Stephens (2013)^1^ reasoned that the correlation between Z-scores for the two phenotypes under the null is the same as the phenotypic correlation, thus “a set of putative null SNPs” were selected, by taking SNPs with |*z*| *<* 2. The same idea was also adopted by later studies^2, 4^ The tool metaCCA^3^ neglected the null effect requirement, as the genetic effect per variant is tiny, and computed the correlation between Z-scores across as many SNPs as possible. However, the Z-cut estimator can generate bias due to its constrain on the summary statistics of the SNPs^7^. LDSC intercept performs better thus was adopted in statistical methods that requires pre-calculated phenotypic correlations^6, 7^, but the intercept collects noise generated by e.g., population substructure, which may also lead to biased estimates of phenotypic correlations^8^. Here, we revisit the correlation between GWAS summary statistics of two phenotypes and propose an alternative approach to select variants for the Z-score correlation estimation strategy. We show that selecting SNPs with low minor allele frequencies (MAFs) can lead to simple and consistent estimation of phenotypic correlations based on multi-SNP Z-score correlations. Via simulations, we show that the “low-MAF” estimator can overcome bias generated by the Z-cut estimator and the LDSC intercept. With higher estimation efficiency, when applied to UK Biobank GWAS results, the low-MAF estimator could discover 30% more significant phenotypic correlations than using the LDSC intercept.

## Methods

We start by deriving a general mathematical form of the correlation between the summary statistics of two phenotypes *y*_1_ and *y*_2_, centred at a zero mean. For a single genetic variant in an association analysis, the model is *y*_*i*_ = *g*_*i*_*β*_*i*_ + *e*_*i*_ (*i* = 1, 2), where *g*_*i*_ is the vector of genotypic values with 0-1-2 coding, and *e*_*i*_ are the residuals. Assuming Hardy-Weinberg equilibrium (HWE), for SNP *j*, we have *g*_*ij*_ ∼ *B*(2, *f*_*j*_), where *f*_*j*_ is the MAF of SNP *j*, and *B*(*n, p*) stands for the binomial distribution with size *n* and success probability *p. g*_1_ and *g*_2_ may differ due to different levels of sample overlap between the two phenotypes. At the single SNP *j* (omitted the subscripts *j* for simplicity),

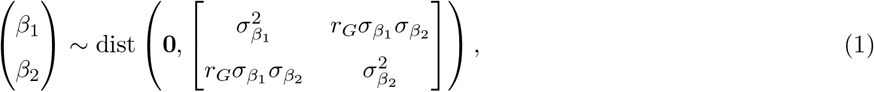

and

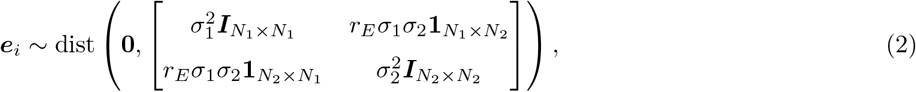

where *r*_*G*_ is the underlying genetic correlation at SNP *j*, and *r*_*E*_ is the residual correlation. In an association study, *r*_*G*_ is un-identifiable at a single SNP. The estimated genetic effects are 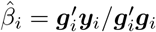, then

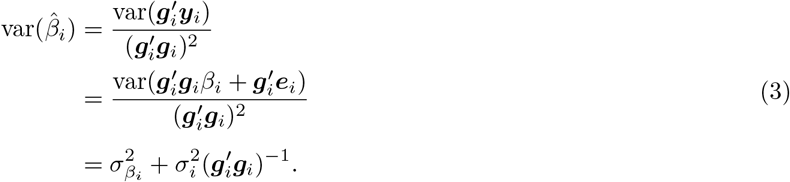

So that

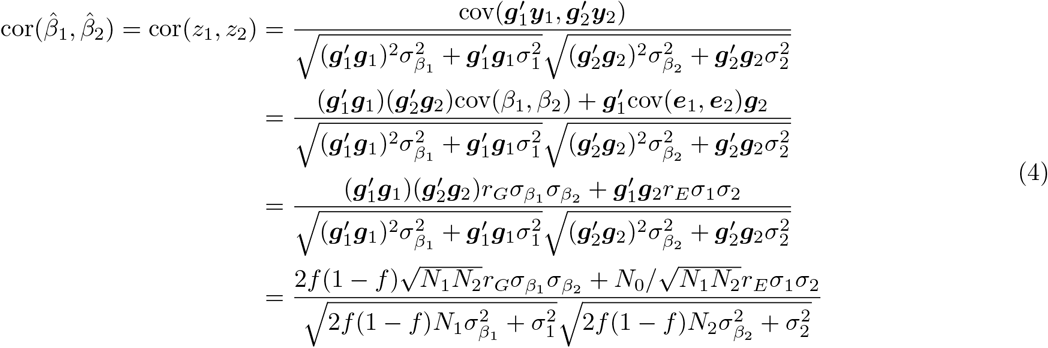

When 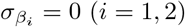, i.e., for any variant with null genetic effect, the above equation simplifies as

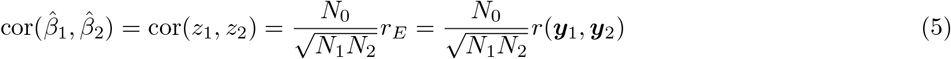

where *r*(*y*_1_, *y*_2_) is the phenotypic correlation based on completely overlapped individual-level data. Particularly, for perfectly overlap samples, i.e., *N*_0_ = *N*_1_ = *N*_2_, we have cor(*z*_1_, *z*_2_) = *r*(*y*_1_, *y*_2_), which is the same as the phenotypic correlation estimator derived by Zhu et al.^2^.

The result suggests that the phenotypic correlation between the two phenotypes *y*_1_ and *y*_2_, subject to a shrinkage factor corresponding to sample overlap, can be estimated by the sample correlation of the summary statistics across any sufficient number of null variants. This leads to a commonly adopted strategy of estimating the phenotypic correlation from summary association statistics by taking a subset with e.g., |*z*_*i*_| *<* 2 (*i* = 1, 2). However, we will show that such thresholding may introduce bias into the correlation estimate.

According to eq. (4), null genetic effect for the variant is a sufficient but not necessary condition for cor(*z*_1_, *z*_2_) to reduce to eq. (5). When *f* = 0, eq. (4) also becomes (5). In practice, the phenotypic correlation can be estimated by the correlation of the summary statistics across a sufficient number of variants with very low minor allele frequencies (MAFs), *regardless* of whether the genetic effects are null. The thresholding on the MAF does not directly introduce a threshold on *β*_*i*_ or *z*_*i*_ so that not prone to bias in the phenotypic correlation estimation.

### Simulation settings

We conducted two sets of simulations to compare the low-MAF estimator with the Z-cut estimator and LDSC intercept, respectively. For the first simulation, we simulated the genotypes of 5,000 independent SNPs in 500 individuals, and the MAFs ranged from 5 × 10^−5^ to 0.5. The genotypes of SNP *j* follow HWE. Two scenarios of phenotypic correlations were evaluated, where in one the phenotypic correlation was set to 0.5 without genetic correlation; and in the other a genetic correlation of 0.5 and a residual correlation of 0.25 were simulated, where the genetic effects across the 5,000 SNPs were extracted from a normal distribution with zero mean. In the simulation, three cutoffs of |*z*| *<* 2, |*z*| *<* 1, and |*z*| *<* 0.5 were evaluated for the Z-cut estimator. Five thresholds of MAFs: 0.5, 0.05, 5 × 10^−3^, 5 × 10^−4^, and 5 × 10^−5^ were evaluated for the low-MAF estimator. The true phenotypic correlations were computed as the Pearson’s correlations of the two vectors of simulated phenotypic values.

For the second simulation, in order to compare with LDSC, we used the real UK Biobank (UKB) genotypes for 336,000 genomic British individuals across the 1,029,876 quality-controlled HapMap3 SNPs selected by the high-definition likelihood (HDL) software^9^. We draw the genetic effects across 10% of the SNPs from a normal distribution with zero mean, so that the phenotypic, genetic, and residuals correlations all had a true value of 0.5. 70,042 SNPs with MAF *<* 5 × 10^−4^ were selected for the low-MAF estimator. Two reference panels were evaluated for LDSC, including the ldsc software inbuilt 1000 Genomes reference and the UKB reference based on the HDL software reference data.

## Results

### The low-MAF estimator corrects the bias of the Z-cut estimator

In the first simulation setting, when no genetic effect was present, namely, every SNP had a null effect, the Z-score correlation estimator based on all the SNPs satisfied eq. (5), resulted in unbiased estimates of the phenotypic correlations. However, constraining the Z-cut estimator on SNPs filtered by Z-score cutoffs generated downward-biased estimates. On the other hand, constraining the low-MAF estimator did not generate bias, regardless of the MAF cutoff (**Fig. 1a**). When genetic correlation was present, the Z-score correlation estimator based on all the SNPs produced inflated estimates, as the common SNPs with large MAFs substantially contributed to the genetic correlation. Same as in the previous scenario, the Z-cut estimator generated downward-biased estimates. With sufficiently low MAF cutoffs, the Z-score correlation maintained as a consistent estimator of the phenotypic correlation (**Fig. 1b**). Also, the estimation efficiency of the low-MAF estimator attained that of the estimator based on observed phenotypic values (**Table 1**).

**Table 1.**
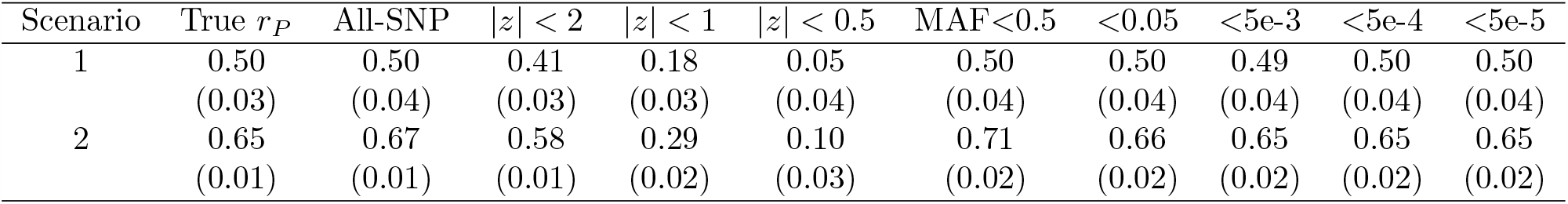
Comparison of the phenotypic correlation estimates by the low-MAF and Z-cut estimators. The results are means and standard deviations (in brackets) summarised from 100 replicates, where in each replicate, 5,000 SNPs were simulated for 500 individuals, and the minor allele frequencies (MAFs) ranged from 5e-5 to 0.5. The true (phenotypic) correlations (*r*_*P*_) were computed as the Pearson’s correlations of the two vectors of simulated phenotypic values. Scenario 1: The two phenotypes had no genetic correlation and a (residual) phenotypic correlation of 0.5; Scenario 2: The two phenotypes had a genetic correlation of 0.5 and a residual correlation of 0.25.

**Fig 1.**
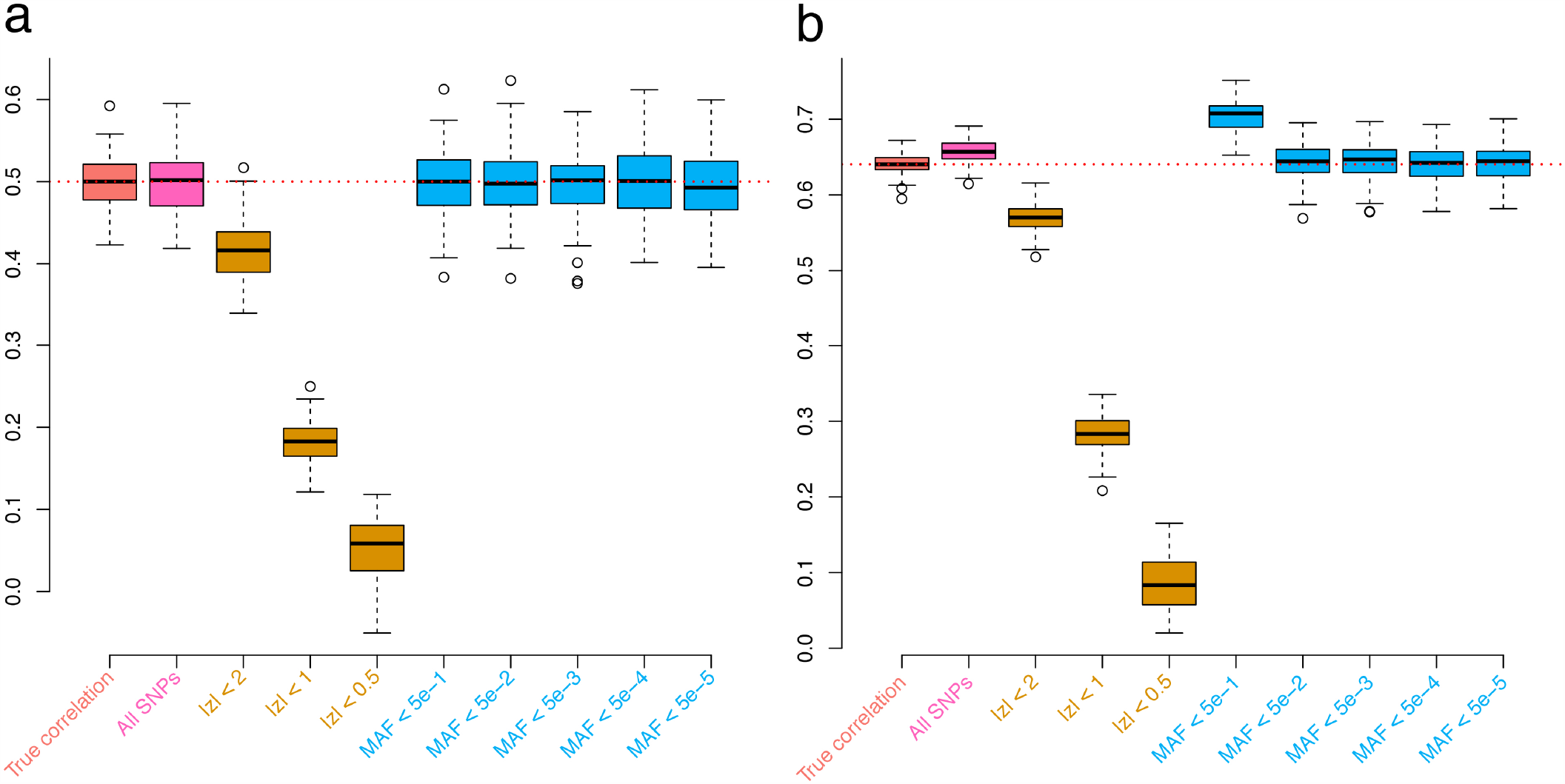
Simulations comparing the Z-cut and low-MAF estimators for phenotypic correlation. The boxplots show the results from 100 replicates, where in each replicate, 5,000 SNPs were simulated for 500 individuals, and the minor allele frequencies (MAFs) ranged from 5e-5 to 0.5. The true (phenotypic) correlations were computed as the Pearson’s correlations of the two vectors of simulated phenotypic values. **a**. The two phenotypes had no genetic correlation and a (residual) phenotypic correlation of 0.5; **b**. The two phenotypes had a genetic correlation of 0.5 and a residual correlation of 0.25.

### The low-MAF estimator corrects the bias of LDSC intercept

For the second simulation, we observed downward bias in the LDSC intercept when the default 1000 Genomes reference was applied (**Fig. 2**). Such a bias was overcome by the UKB reference, nevertheless, the estimates were slightly inflated possibly due to the population substructure in the UKB genomic British individuals^8^. These biases were all absent when applying the low-MAF estimator for the phenotypic correlation. Furthermore, the low-MAF estimator had a substantially higher estimation efficiency than the LDSC intercept, as if the sample size was 10 times larger (**Table 2**).

**Table 2.** Comparison of the phenotypic correlation estimates by the low-MAF estimator and LDSC intercept. The results were summarised from 100 replicates, where in each replicate, two phenotypes were simulated for 336,000 genomic British individuals. The true phenotypic, genetic, and residual correlations were all set to 0.5. The low-MAF estimates were based on 70,042 SNPs with MAF *<* 5e-4. 1kG ref: LD scores calculated based on the 1000 Genomes reference panel; UKB ref: LD scores calculated based on the UK Biobank reference panel.

|  | Low-MAF | LDSC (1kG) | LDSC (UKB) |
| --- | --- | --- | --- |
| Minimum | 0.4865 | 0.4447 | 0.4656 |
| 25% Quantile | 0.4964 | 0.4719 | 0.4927 |
| Median | 0.4996 | 0.4887 | 0.5034 |
| Mean | 0.5001 | 0.4871 | 0.5040 |
| 75% Quantile | 0.5035 | 0.4997 | 0.5166 |
| Maximum | 0.5205 | 0.5391 | 0.5498 |
| Variance | 3.603e-05 | 3.626e-04 | 3.022e-04 |

**Fig 2.**
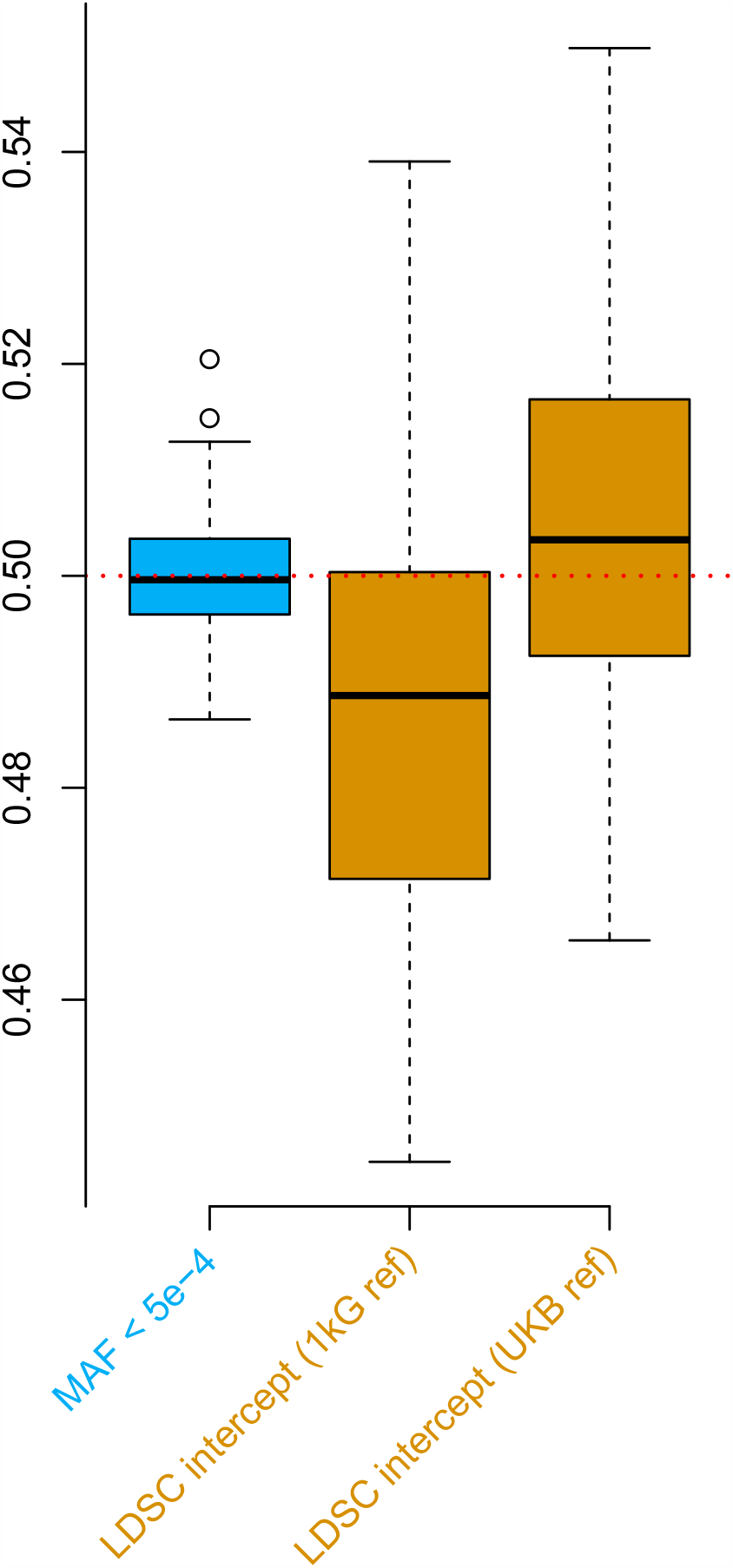
Simulations comparing the low-MAF estimator and LD score regression (LDSC) intercept using the UK Biobank genotype data. The boxplots show the results from 100 replicates, where in each replicate, two phenotypes were simulated for 336,000 genomic British individuals. The true phenotypic, genetic, and residual correlations were all set to 0.5. The low-MAF estimates were based on 70,042 SNPs with MAF *<* 5 × 10^−4^. 1kG ref: LD scores calculated based on the 1000 Genomes reference panel; UKB ref: LD scores calculated based on the UK Biobank reference panel.

### Example

We selected the same 30 GWASed phenotypes used by Ning et al.’s in genetic correlation estimation^9^, as a real data example to compare the low-MAF estimator to LDSC intercept in the estimation of the phenotypic correlations (**Fig. 3**). The low-MAF estimates were based on 70,042 SNPs with MAF *<* 5 × 10^−4^, and the LD scores were calculated based on the 1000 Genomes reference panel (default). At a 5% Bonferroni-corrected p-value threshold for 435 pairs of traits, the low-MAF method discovered 223 significant phenotypic correlations, and LDSC intercept discovered 171. Among these, 61 phenotypic correlations were only significant in the low-MAF method, versus 9 only significant using the LDSC intercept.

**Fig 3.**
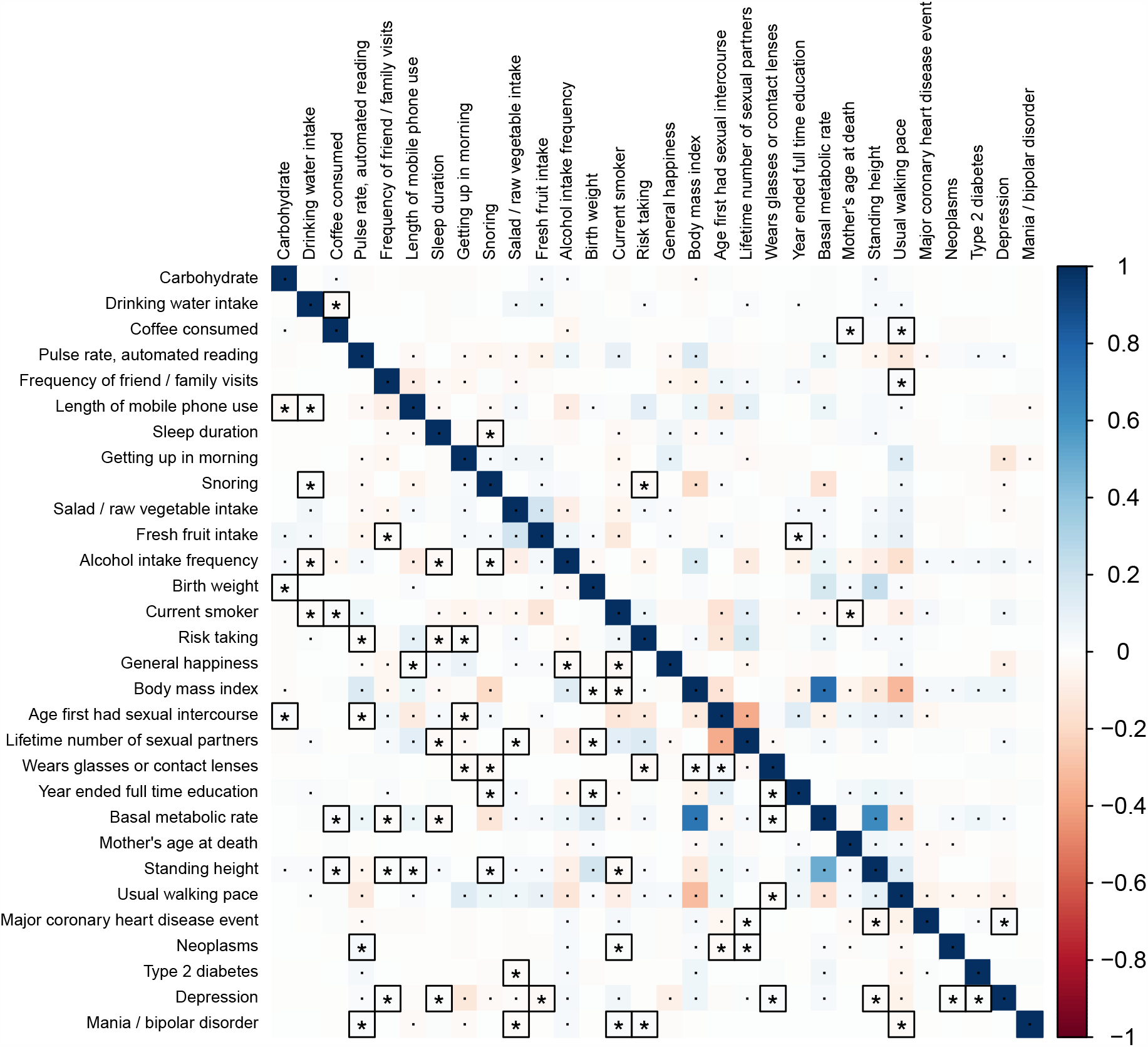
Phenotypic correlations across 30 UK Biobank traits using the low-MAF estimator (lower triangle) and LD score regression (LDSC) intercept (upper triangle). The default 1000 Genomes reference panel was used in LDSC. Bonferroni-corrected significant correlations with *P <* 0.05*/*435 are marked with asterisks or dots, where those correlations that are only significant using one of the two methods are marked with asterisks and squares.

## Discussion

We have proposed the low-MAF estimator of phenotypic correlations based on GWAS summary statistics, as an improvement of the Z-score correlation strategy based on all SNPs or SNPs that pass a particular Z-score cutoff. The estimator overcomes the bias generated when thresholding on summary association statistics and even that generated in the bivariate LDSC intercept. We suggest the use of the low-MAF phenotypic correlation estimator in future practice. The more consistent and efficient estimation can improve our understanding of connections across human complex traits and diseases.

Although the low-MAF method also introduces a filter on the tested SNPs, it is a threshold-free technique for the genetic effect parameter. Thus, the low-MAF estimator does not constrain the estimated genetic effects of selected SNPs, equivalent to sampling a set of null effect SNPs from the genome. This explains why “putative null effect” SNPs with e.g., |*z*| *<* 2 generate bias whereas the low-MAF estimator does not. Even if all the SNPs are null, some of them will generate z-score with |*z*| *>* 2 due to randomness. Removing them would lead to bias.

As the low-MAF estimator is equivalent to sampling a set of null effect SNPs from the genome, the resulted phenotypic correlation estimates are close to those estimated using individual-level phenotypic data. In the real UKB genotype data simulation, we showed that the LDSC intercept could not produce consistent estimates of the phenotypic correlation due to population substructure. Such a complication in LDSC was overcome by the low-MAF estimator, because although the GWAS summary statistics were used, the estimator approximates observed phenotypic correlation and is irrelevant to genetic data structure. Namely, the genotypic data are treated as nuisance in the low-MAF estimator.

For binary phenotypes, an advantage of summary-statistics-based estimators, such as the low-MAF estimator, is that it estimates the underlying phenotypic correlations on the liability scale. The liabilities follow an unobservable logistic distribution therefore the estimates are not the same as the observed phenotypic correlations directly computed using the 0-1 outcome data. The phenotypic correlation estimates on the liability scale is mathematically easier to interpret and can be transformed into odds ratios from logistic regressions.

## Data availability

The individual-level genotype and phenotype data are available by application from the UKBB (http://www.ukbiobank.ac.uk/). The UKBB GWAS summary statistics by the Neale laboratory can be obtained from http://www.nealelab.is/uk-biobank/.

## Code availability

HDL: https://github.com/zhenin/HDL; ldsc: https://github.com/bulik/ldsc.

## Acknowledgements

X.S. was in receipt of a Swedish Research Council (VR) Starting Grant (No. 2017-02543). We thank Edinburgh Compute and Data Facility at the University of Edinburgh and National Supercomputer Center at Sun Yat-sen University for providing high-performance computing resources.

## Author contributions

X.S. initiated and coordinated the study. T.L. and Z.N. contributed to data analysis. X.S. drafted the manuscript. All authors contributed to manuscript writing and gave final approval to publish.

## Ethics declarations

The authors declare no competing interests.

## References

[1] Stephens, M. A unified framework for association analysis with multiple related phenotypes. PLOS ONE 8(7), e65245 (2013).

[2] Zhu, X., Feng, T., Tayo, B. O., Liang, J., Young, J. H., Franceschini, N., Smith, J. A., Yanek, L. R., Sun, Y. V., Edwards, T. L., Chen, W., Nalls, M., Fox, E., Sale, M., Bottinger, E., Rotimi, C., Consortium, C. B. P., Liu, Y., McKnight, B., Liu, K., Arnett, D. K., Chakravati, A., Cooper, R. S., and Redline, S. Meta-analysis of correlated traits via summary statistics from GWASs with an application in hypertension. American journal of human genetics 96(1), 21–36 (2015).

[3] Cichonska, A., Rousu, J., Marttinen, P., Kangas, A. J., Soininen, P., Lehtimäki, T., Raitakari, O. T., Järvelin, M.-R., Salomaa, V., Ala-Korpela, M., Ripatti, S., and Pirinen, M. metaCCA: summary statistics-based multivariate meta-analysis of genome-wide association studies using canonical correlation analysis. Bioinformatics 32(13), 1981–1989, jul (2016).

[4] Shen, X., Klarić, L., Sharapov, S., Mangino, M., Ning, Z., Wu, D., Trbojević-Akmačić, I., Pučić-Baković, M., Rudan, I., Polašek, O., Hayward, C., Spector, T. D., Wilson, J. F., Lauc, G., and Aulchenko, Y. S. Multivariate discovery and replication of five novel loci associated with Immunoglobulin G N-glycosylation. Nature Communications 8(1), 447, dec (2017).

[5] Bulik-Sullivan, B., Finucane, H. K., Anttila, V., Gusev, A., Day, F. R., Loh, P.-R., Consortium, R., Consortium, P. G.G. C. f. A. N. o. t. W. T. C. C. C., Duncan, L., Perry, J. R. B., Patterson, N., Robinson, E. B., Daly, M. J., Price, A. L., and Neale, B. M. An atlas of genetic correlations across human diseases and traits. Nature genetics 47(11), 1236–1241 (2015).

[6] Turley, P., Walters, R. K., Maghzian, O., Okbay, A., Lee, J. J., Fontana, M. A., Nguyen-Viet, T. A., Wedow, R., Zacher, M., Furlotte, N. A., Magnusson, P., Oskarsson, S., Johannesson, M., Visscher, P. M., Laibson, D., Cesarini, D., Neale, B. M., and Benjamin, D. J. Multi-trait analysis of genome-wide association summary statistics using MTAG. Nature Genetics 50(2), 229–237, feb (2018).

[7] Zheng, J., Richardson, T. G., Millard, L. A. C., Hemani, G., Elsworth, B. L., Raistrick, C. A., Vilhjalmsson, B., Neale, B. M., Haycock, P. C., Smith, G. D., and Gaunt, T. R. PhenoSpD: an integrated toolkit for phenotypic correlation estimation and multiple testing correction using GWAS summary statistics. GigaScience 7(8), 1–10, aug (2018).

[8] Yengo, L., Yang, J., and Visscher, P. M. Expectation of the intercept from bivariate LD score regression in the presence of population stratification. bioRxiv 0(0), 0–0 (2018).

[9] Ning, Z., Pawitan, Y., and Shen, X. High-definition likelihood inference of genetic correlations across human complex traits.Nature Genetics 52(8), 859–864, aug (2020).

